## Supplemental Data for "Homothorax Controls a Binary Rhodopsin Switch in *Drosophila* Ocelli"

**Figure S1**

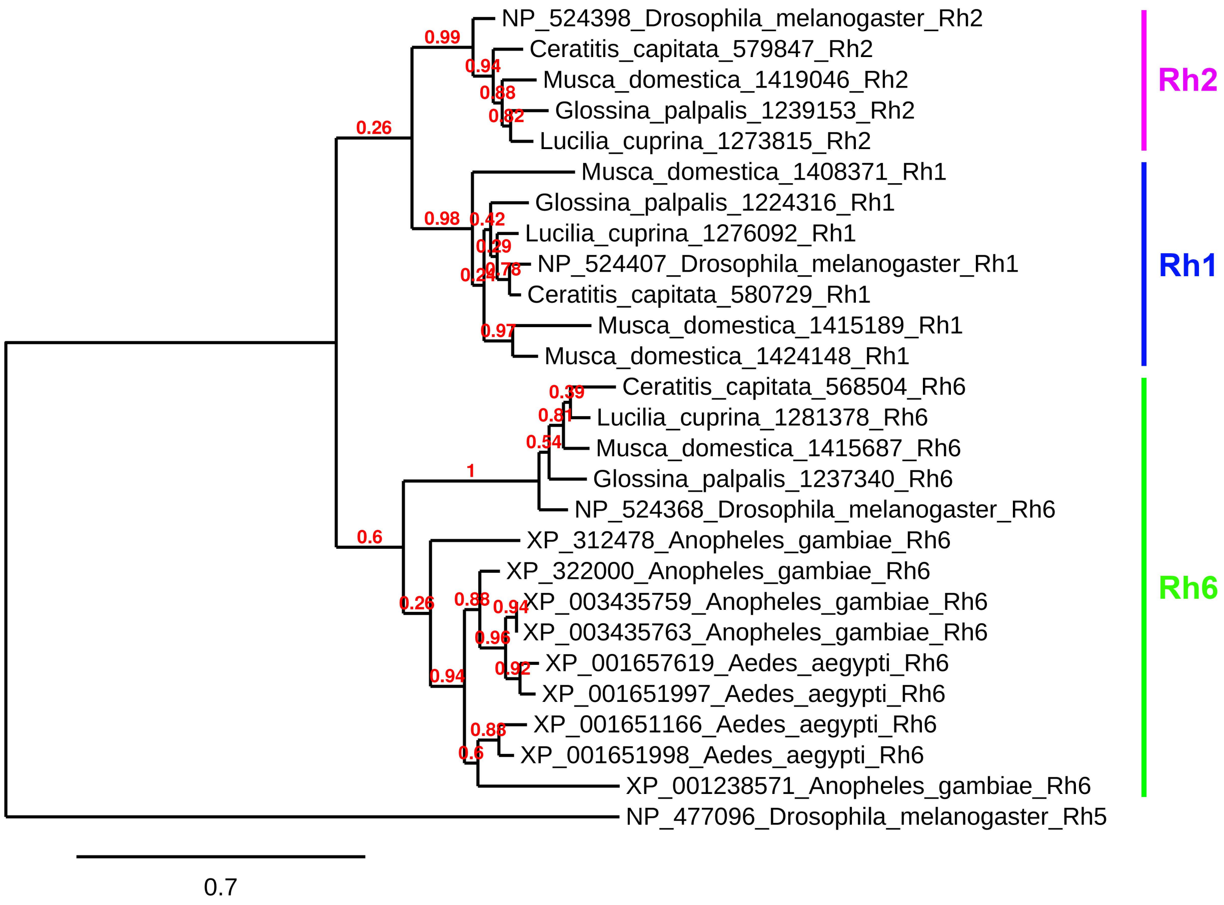

NP_524398_Drosophila_melanogaster_Rh2
Ceratitis_capitata_579847_Rh2
Musca_domestica_1419046_Rh2
Glossina_palpalis_1239153_Rh2
Lucilia_cuprina_1273815_Rh2
Musca_domestica_1408371_Rh1
Glossina_palpalis_1224316_Rh1
Lucilia_cuprina_1276092_Rh1
NP_524407_Drosophila_melanogaster_Rh1
Ceratitis_capitata_580729_Rh1
Musca_domestica_1415189_Rh1
Musca_domestica_1424148_Rh1
Ceratitis_capitata_568504_Rh6
Lucilia_cuprina_1281378_Rh6
Musca_domestica_1415687_Rh6
Glossina_palpalis_1237340_Rh6
NP_524368_Drosophila_melanogaster_Rh6
XP_312478_Anopheles_gambiae_Rh6
XP_322000_Anopheles_gambiae_Rh6
XP_003435759_Anopheles_gambiae_Rh6
XP_003435763_Anopheles_gambiae_Rh6
XP_001657619_Aedes_aegypti_Rh6
XP_001651997_Aedes_aegypti_Rh6
XP_001651166_Aedes_aegypti_Rh6
XP_001651998_Aedes_aegypti_Rh6
XP_001238571_Anopheles_gambiae_Rh6
NP_477096_Drosophila_melanogaster_Rh5
Rh2
Rh1
Rh6
0.99
0.94
0.88
0.82
0.26
0.98
0.42
0.29
0.78
0.24
0.97
0.39
0.81
0.54
1
0.6
0.26
0.88
0.94
0.96
0.92
0.94
0.88
0.6
0.7

### Figure S2

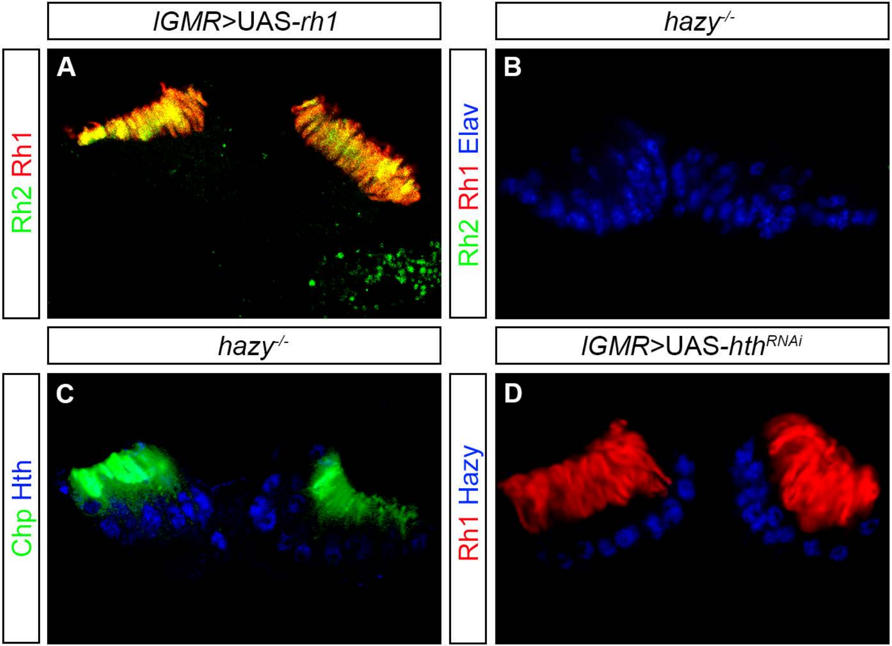

lGMR>UAS-rh1
hazy-/-
A
Rh2 Rh1
B
Rh2 Rh1 Elav
hazy-/-
lGMR>UAS-hthRNAi
C
Chp Hth
D
Rh1 Hazy

### Figure S3

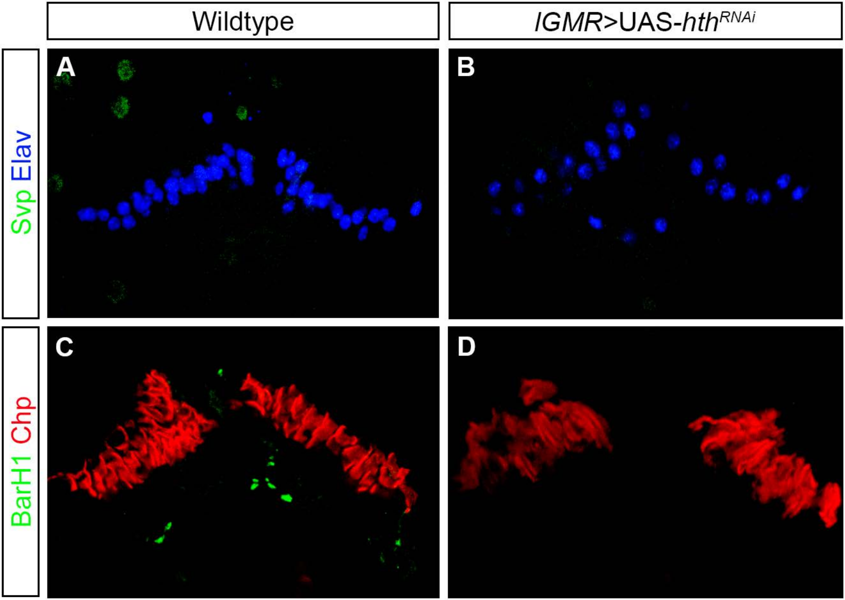

**Figure S4**

### rhodopsin 1 promoter

```
                                                                                  hth1
Dmel    1 CTGGAGACTCAAGAATAATACTCGGCCAGAACCCCCAGTGGAAACCCTTGAAATGC-------------CTTTAACTATTGACATTGGCCCATTGCGATG  87
Dsim    1 CTGGA--CACAAGACTAATGCTCGGCCAGAACCCCCCGTGGAAACCCTTGAAATGC-------------CTTTAACTATTGACATTGGCCCATTGCGATG  85
Dsec    1 CTAGG--CACAAGCCTAATGCTCGGCCAGAACCCCCCGTGGAAACCCTTGAAATGC-------------CTTTAACTATTGACATTGGCCCATTGCGATG  85
Dyak    1 CTCGG--GAGAAGACTAATGCTCGGCCAGAACCCTC-TTGGAAACCCTTGAAATGC-------------CTTTAACTATTGACATTGGCCCATTGCGATG  84
Dere    1 CTCGG--CAGAAGACTAATGCTCGGCCAGAACCCGTCTTGGAATCCGTTGAAATGC-------------CTTTAACTATTGACATTGGCCCATTGCGATG  85
Dana    1 ------------------------------GCCGTTCTCGGAAACCATTGAAATGTCTTTGGCGAAAGACTTTAGCTATTGACATTGGCCCACGGCGATG  70
Dper    1 GACGAGACCCATAGATACCCATAATCGAGAGGTGCTGAAAGAAGCCTCTGAAATGTC---------------TAACTATTGACATTGGCCCATTGC----  81
Dpse    1 GACGAGACCCATGGATACCCATAATCGAGAGGTGCTGAAAGAAGCCTCTGAAATGTC---------------TAACTATTGACATTGGCCCATTGC----  81
Dwil    1 TTAAATCGCTGACTTTCCACGCCCCCAATTTTGCACAATGTCAAAAGCTTGAAG--CCTTTA-----------AAGTATTGACATTGATTAGCTTTGTTG  87
Dgri    1 GTCAGCGCATATGTTGTTGCCACGCCTGCCGGTGCAGCGTGGAAACCTGTAAAGT-------ACAAGAACTATAACTATTGACATTGGACAGCGTTGTTC  93
Dmoj    1 CAAACGTTGGCCCTCTGCCAGCCCAGCGAATCTGGAGCTGGGAAACCTTTGAAGT--------------ACATAACTATTGACATTGGATGTCGTCGCGT  86
Dvir    1 CAAGTGTTGGCCCGTTGCCAGCCGGGCACATCTCGAGCTTGGAAACCTTTGAAGTACCTTTAACAAGAACCATAACTATTGACATTGGGCTAGCCAGTTG 100

                                                                                   hth2
Dmel   88 TGCG-----------------------------------CTTGCTTTCGTTATGGATATTATGATTATAAGCC-AAAA-GCTGTCGACACTTTCCTCTGC 150
Dsim   86 TGCG-----------------------------------CTTGCTTTCGTTATGGCTATTATGATTATAAGCC-AAAA-GCTGTCGACACTTTCCTCTGC 148
Dsec   86 TGCG-----------------------------------CCTGCTTTCGTTATGGCTATTATGATTATAAGCC-AAAA-GCTGTCGACACTTTCTTCTGC 148
Dyak   85 CGCG-----------------------------------CTTGCTTTTCTTATGGCTATTATGATTATAAGCC-AAAA-GCTGTCGACACTTTCCTCTGC 147
Dere   86 TGCG----------------------------ATGTGCGCTTGCTTTTGTTATGGCTATTATGATTATAAGCC-GAAA-GCTGTCGACACTTTCCTCTGC 155
Dana   71 TGCA-----------------------------------CTGGATTTTGTTATGACTATTATGATTATAAGCCAGAAA-GCTGTCGACATTTTTCTCCTC 134
Dper   82 -------------------------------------------------AATGGGCTATTATGATTATAAGCC-AAAA-GCTGTCTGCACTTTTGTAGCA 130
Dpse   82 -------------------------------------------------AATGGGCTATTATGATTATAAGCC-AAAA-GCTGTCTGCACTTTTGTAGCA 130
Dwil   88 AACAAACAGAAGAATATTTCACTCATTTTTGCCATTTGTTATGGTTCTTATATCTGGTGTATGGTTATAAGCCCAAAAAGCTGTTAACACTTTCCTTTTG 187
Dgri   94 GTCAAATTGTATT-----------TTATATTTTCTTGTGCATTTTGTTTGGTTTAACGTTTATGTTATAAGCC-CTTA-GCTGTTAACGCATATA----- 175
Dmoj   87 CATCGTTGCAATA-----------TTATATTTGTTATATTT---------GTTTAA-GTTTACGTTATAAGCC-AAAG-GCTGTTAACGCCTT------- 156
Dvir  101 TCGCGTCATCGTT----GGCATTATTATATTTTATATTTTGTGAAATATTTCTTTTGGTTTGTTTTATAAGCC-AAAA-GCTGTTAACGCCTT------- 187

                                                                             RCSI
Dmel  151 ACATTGCAGACA---------------TATTTTTAGG-------------------------CTGGCCTAATTGAATTTCAAGAGCA------GGGGTT- 203
Dsim  149 ACACTGCACACA---------------TATTTTTAGG-------------------------CTAGCCTAATTGAATTTCAAGAGCA------GGGGTT- 201
Dsec  149 ACATTGCACACA---------------TATTTTTAGG-------------------------CTAGCCTAATTGAATTTCAAGAGCA------GGGGTT- 201
Dyak  148 ACACTGCACACA---------------TATTTTTAGG-------------------------CTGGCCTAATTGAATTTCAAGACTA------GGGGTT- 200
Dere  156 ACACTGCACACA---------------TATTTTTAGA-------------------------CTGGCCTAATTGAATTTCAAGAGCA------TGGGTT- 208
Dana  135 AACGCTTTCTTCACATTGCTGGAGGTGTATTTTTAGGCC---------------------------CCTAATTGAATTTGAATTTC-------TGGGTT- 199
Dper  131 TGTACGTATGTATA-------------TATTTTTAGG--------------TGCACTGGCGTCTTGGCTAATTGAATTTGAACCCTACCAGAGGGGGTTC 203
Dpse  131 TGTACGTATGTATA-------------TATTTTTAGG--------------TGCACTGGCGTCTTGGCTAATTGAATTTGAACCCTACCAGAGGGGGTTC 203
Dwil  188 CATT-----------------------TATTTTTAGGCCTCCAGCGTCAAGTGCATTTT------GTCTAATTGAATTTA-------------------- 238
Dgri  176 ---------------------------TATTTTTAGGCCTTTG------AGTGCAGCAA--------CTAATTCAATTTGTA------------------ 216
Dmoj  157 ---------------------------TATTTTTAGGCCTCCG------AGTGCAAGAA--------CTAATTGAATTTGTTTGG--------------- 200
Dvir  188 ---------------------------TATTTTTAGGCCTTTG------AGTGCAAGAA--------CTAATTGAATTTGGTTTG--------------- 231

                                                 exon1 5'UTR
Dmel  204 ----ACACGAGCACCGCGAAAACTATAAAAGCCATGCGGCAGAATGCAGACATTGCAGGTTTCCAACGACCAATCGCCGCGACTAGTCCGCCCCAGTGAA 299
Dsim  202 ----ACACGAGCACCGGGAAAACTATAAAAGCCACGCGGCAGAATGCAGACAATGCAGGTTTCCAACGACCAATCGCCGCGAATAGTCCGCCCCAGTGAA 297
Dsec  202 ----ACACGAGCACCGGGAAAACTATAAAAGCCACGCGGCAGAATGCAGACAATGCAGGTTTCCAACGACCAATCGCCGCGAATAGTCCGCCCCAGTGAA 297
Dyak  201 ----ACACGAGCGCCACGAAAACTATAAAAGCCAGGGTTCAGAATGCAGACAATGCAGGTTTCCAACGACCAATCGCCGCGAATAGTCCGCCC-AGTGAA 295
Dere  209 ----ACAGAGCACCGCGGAAAACTATAAAAGCCAGGGTGCAGAATGCAGACAATGCAGGTTTCCAACGACCAATCGCCGCGAACAGTCCGCCCCAGTGAA 304
Dana  200 ----CTGCAATGGCGGGAAAAACTATAAAAGCCAGGTCAAGAAGTGCAGTCGATGCAGGTTTCCAACGACCAATCGCCGCGAATAGTCCACCCCAGTGAA 295
Dper  204 TTTTAAGAATCTGCAAGGAAAACTATAAAAGCCCCAGAGTCACCAGCGGACGTTGCAGGTTTCGAACGACCAATCGCCGCGAATAGTCCGCTCCAGTGAA 303
Dpse  204 TTTTAAGAATCTGCAAGGAAAACTATAAAAGCCCCAGAGCCACCAGCGGACGTTGCAGGTTTCGAACGACCAATCGCCGCGAATAGTCCGCTCCAGTGAA 303
Dwil  239 -----------GTGAGGAAAAACTATAAAAACCACACAACATTTGTCGAACGATGCAGGTTTCCAACGACCAATCGCCGCGAATAGTCCGCTCCAGTGAA 327
Dgri  217 ----CTTTAAACGTTGGGAAAACTATAAAAGCGACGTCTGAGCCAGGCACAGACGCAGGTTTCCAACGACCAATCGCCGCGAATAGTCCGCTTCAGTGAA 312
Dmoj  201 ----ACATGGGCCAAGACAAGACTATAAAAGCAACGCTTGAGCCAGGCACAGACGCAGGTTTCCAACGACCAATCGCCGCGAATAGTCCGCTCCAGTGAA 296
Dvir  232 --------TTTAGGAAAATAAACTATAAAAGCGGCGTCCCAGCCAGGCACAGACGCAGGTTTCAAAAAACCAATCGCCGCGAATAGTCCGCTCCAGTGAA 323

Dmel  300 ATATTCAGAATCCAGGAACCCTT------TATGTAAAAAGTGTTAGAAATATTGTTAGTG------------------------AATTTGCA-GCTTTTT 368
Dsim  298 ATATTCAGAATCCAGGAACCCTT------TATGTAAAAAGTGTTAGAAATATTGTTAGTG------------------------AATTTGCA-GCTTTTT 366
Dsec  298 ATATTCAGAATCCAGGAACCCTT------TATGTAAAAAGTGTTAGAAATATTGTTAGTG------------------------AATTTGCA-GCTTTTT 366
Dyak  296 ATATTCAGAATCCAGGAACCCTT------TATGTAAAAAGTGTTAGAAATATTGTTAGTG------------------------AATTTGCT-GCTTTTT 364
Dere  305 ATATTCAGAATCCAGGAACCCTT------TATGTAAAAAGTGTTAGAAATATTGTTAGTG------------------------AATTTGCA-GCTTTTT 373
Dana  296 ATATTCAGAGTTCAGGAGTCCTT------TATGTAAAAAGTGTTTGAAATATTAATAGTG------------------------AATTTGCA-GCTTTTT 364
Dper  304 ATATTCAAAATAAATTTAGGGAATCCTTTAATGTAAAAAGTTTTTGAAAAATATCTAACCG-----------------------AATTTGCAAGCTTTTT 380
Dpse  304 ATATTCAAAATAAATTTAGGGAATCCTTTAATGTAAAAAGTTTTTGAAAAATATCTGACCG-----------------------AATTTGCAAGCTTTTT 380
Dwil  328 ATATTCATAATAATTAACAGATT--CCTTTATGTAAAGAAAGTTTTGTTTTGAGAT-------------------CCAAGTGAATTTTTGC-AGCTTTGT 405
Dgri  313 ATATTCAGGGAATAACTAATTTT-------ATGTAAAAAGTGTT-GAAATTTAAAAAAAT---------AAAAAATAAAACGTTTATTTGC-TGCTTTTT 394
Dmoj  297 ATAATATTCAGGGAAATTACAAAATCT---ATGTAAAAAGTGTTCGAAATTTAGAGAGAT-----------------TAAAACTAATTTGC-AGCTTTTT 375
Dvir  324 AAACATTCAGGGAAAGATT------------TGTAAAAAGTGTTAGAGATTTAAAAATTATTACAAAAAAAAAAAATAAATATTAATTTGC-AGCTTTTT 410

                                                                                       Met            intron
Dmel  369 ATGTAGACAGTGTGA-----------TAT-AGGCGGGATATAGTGACGCAGCCAGTAACCAAAA-----------------CACA-ATG-GAG-AG-GTA 435
Dsim  367 ATGTAGACAGTGTGA-----------TAT-AGGCGGGATATAGTGACGCAGCCAGTAATCAAAA-----------------CACA-ATG-GAG-AG-GTA 433
Dsec  367 ATGTAGACAGTGTGA-----------TAT-AGGCGGGATATAGTGACGCAGCCAGTAATCAAAA-----------------CACA-ATG-GAG-AG-GTA 433
Dyak  365 ATGTAGACAGTGTGA-----------TAT-AGGCGGGATATAGTGACGCAGCCAGTAACCAAAA-----------------CACA-ATG-GAG-AG-GTA 431
Dere  374 ATGTAGACAGTGTGA-----------TAT-AGGCGGGATATAGTGACGCAGCCAGTAACCAAAA-----------------CACA-ATG-GAG-AG-GTA 440
Dana  365 ATGTAGACAGTGTGA-----------TAC-AGGCGGGATATAGTGACGCAGCCAGAAACTTAGA-----------------CACA-ATG-GAG-AG-GTA 431
Dper  381 ATGTAGAAAGAGTGA-------GTTTTAT-AGGCGGGATATAGTGACACACAAATAACTGGGAAAAATAAAAT--------CACA-ATG-GAT-AG-GTA 460
Dpse  381 ATGTAGAAAGAGTGA-------GATTTAT-AGGCGGGCTATAGTGACACACAAATAACTGGGAAAAATAAAAT--------CACA-ATG-GAT-AG-GTA 460
Dwil  406 ATGTAGAAAAGATTGTGTTAAGAATTTAT-AGACGGGATATAGTGTAAATACAAAAAAAAAAAACCACCAAAAACATTTATTAAA-ATG-GAA-AG-GTA 500
Dgri  395 ATGTAAAACGCAGAAGTTAGTGAAAGCAAAAGCCGGGATATAGTGCTAATACCAATACGAGATT-----------------CGAA-ATG-GAG-AG-GTA 473
Dmoj  376 ATGTAAAACGCAGAAGTTAGTGAAAACTTAAGCCGGGATCTAGCGGATACACAAATT------------------------CACA-ATG-GAA-AG-GTA 447
Dvir  411 ATGTAAGTCGCAGAAGCTAGTGAAAGCTAAAGCCGGGATATAGTGCAAAAGCGGATATTTAAAATT---------------CAAA-ATG-GAG-AG-GTA 491
```

### Figure S5

#### rhodopsin 2 promoter

Last exon CG14297

```
Dmel   1 TCG-TCG-ACC-GAC-CCA-TCC-AAC-TCG-AAG-ACC-TCG-ATG-CCA-TCC-TGC-AGC-CAA-GGA-GTG-CAG-CGA-AAA-CTG-TGC-CAG-  75
Dsim   1 TCA-TCG-ACC-GAC-CCA-TCC-AAC-TCG-AAG-ACC-TCG-ATG-CCA-TCG-TGC-AGC-CAA-GGA-GTG-CAG-CGA-AAA-CTG-TGC-CAC-  75
Dsec   1 TCA-TCG-ACC-GAC-CCA-TCC-AAC-TCG-AAG-ACA-TCG-ATG-CCA-TCG-TAC-AGC-CAA-GGA-GTG-CAG-CGA-AAA-CTG-TGC-CAC-  75
Dyak   1 TCG-TCG-ACC-GAC-CCA-TCC-AAT-TCG-AAG-ACC-TCG-ATG-CCG-TCC-TGC-AGC-CAA-GGA-GTA-CAG-CGA-AAA-CTG-TGC-CAG-  75
Dere   1 TCG-TCG-ACC-GAT-CCC-TCC-AAT-TCG-AAG-ACC-TCG-ATG-CCG-TCC-TGC-AGC-CAA-GGA-GTG-CAG-CGA-AAA-CTG-TGC-CAG-  75
Dana   1 TAC-AAT-ACT-AGC-AAG-ATG-GAC-CAC-AAC-ATG-CAG-TGC-AGC-AAA-AAC-TCT-CGT-GGC-ATG-CAG-CGA-AAA-CTG-TGC-CAA-  75
Dpse   1 ACA-AAG-GTC-TTA-GCC-ACG-CCA-GCG-CCC-AAG-TGC-CAC-GAA-AAC-TCA-TCC-AAC-TGC-CGC-CAA-AGG-AAA-CTG-TGC-CTG-  75
Dper   1 ACA-AAG-GTC-TTA-GCC-ACG-CCA-GCG-CCC-AAG-TAC-CAC-GAA-CAC-ACA-TCC-AAC-TGC-CGC-CAA-AGG-AAA-CTG-TGC-CTC-  75
Dwil   1 CTA-AGG-CTA-TTG-CCA-GTG-CAA-GAC-AAA-GAA-CTG-GAC-ACA-AAT-GCA-ACC-CAG-GGA-AGG-CAG-AGA-AAA-CTT-TGT-CAG-  75
Dmoj   1 TCG-AAA-AAC-TCT-CTA-TCA-CCG-TGC-TCA-GTG-TGT-TCC-ATA-GGC-ATG-CCC-CAG-TCG-AAT-AGA-AAG-AAA-CTA-TGC-CAC-  75
Dgri   1 CCC-TCA-AGG-AGC-TCC-TTC-AAC-ATT-TCA-TGT-TCA-TTA-TGT-TCG-GTG-CCT-AAG-TCA-CAT-CAG-AAA-AAG-CTC-TGC-CAA-  75
Dvir   1 AAG-TCC-TCT-TTA-ATC-GAA-CCG-TGC-TCG-GTG-TGC-TCA-ATC-AGT-GCG-CCC-ATG-TCG-AAT-AAA-AAG-AAA-CTT-TGC-CAG-  75

                 hth1                                * hth2
Dmel  76 AAA-TGT-GGA-CAG-AAA-TTT-CTG-GCT-GCC-CTT-TAGTGTCAAGAACTAATATTCAAAGGCGGCAAATTAACCTTGCGCTTA------------ 153
Dsim  76 AAA-TGC-GGA-CAG-AAA-TTT-CTG-GCT-GCC-CTT-TAGTGTCACTAACTAATAATCAAAGGCGGCAAATTAACCTTGCGCTAA------------ 153
Dsec  76 AAA-TGC-GGA-CAG-AAA-TTT-CTG-GCT-GCC-CTT-TAGTGTCACTAACTAATATTCAAAGGCGGCAAATTAACCTTGCGCTAA------------ 153
Dyak  76 AAA-TGC-GGA-CAG-AAG-TTT-CTG-GCT-GCC-CTT-TAGTGTCACTGACTAATATTCAAAGGCG---AATCAATTTTTCTCTAA------------ 150
Dere  76 AAA-TGC-GGA-CAG-AAA-TTT-CTG-GCT-GCC-CTT-TAGTGCCACTGACTAATATTCAAAGGCG---ATTCAACCTTGCACCAA------------ 150
Dana  76 AAG-TGC-GGA-CAA-AAA-TTC-CTT-GCA-GCG-CTT-TGA-ACCTTCGGTGAATATT----GGTT---GACTAGTTTTATGCCAA--AACACAAAGA 155
Dpse  76 CAA-TGC-GGA-AAG-AAA-TTT-TTG-GAC-TCG-CTC-TAG------------------------------------------------------------ 108
Dper  76 CAA-TGC-GGA-AAG-AAA-TTC-TTG-GAC-TCG-CTC-TAG------------------------------------------------------------ 108
Dwil  76 AAT-TGT-AAA-GAA-AAA-TTT-ATG-GCA-GCT-CTG-TGA-----------------------GGAG--AAAG-AATCATAGCCCAT----------- 129
Dmoj  76 AAG-TGT-GGC-CAG-AAA-TTC-TTG-GCT-CAG-CTC-TGA-----------------------------------GCGCCAGCTTCGAATAGACTGTTG 132
Dgri  76 AAG-TGT-GGC-CAG-AAA-TTC-TTG-GCA-AAG-CTC-TAA-----------------------------------------ATAGACGC-AAAACACAAATG 128
Dvir  76 AAA-TGT-GGC-CAG-AAG-TTC-TTA-GCC-AAG-CTC-TGAGCACACACACACACACACACACACACAGACACGCACAGC---AGATGACAAAATG 162

                                                                                                    hth3
Dmel 154 ----------------------------GATTAT----------------GTAGCGTATGAAATTAA-ATAAAATTGACTTTTATCTGGCAGCTGTCC 206
Dsim 154 ----------------------------GATTAT----------------GTAGCGTATGAAATAAA-ATAAAATTGACTTTTATCCGGCAGCTGTCC 206
Dsec 154 ----------------------------GATTAT----------------GTAGCGTATGAAATAAA-ATAAAATTGACTTTTATCCGGCAGCTGTCC 206
Dyak 151 ----------------------------GATTAC----------------GTAGCGTATGAAATAAA-ATGAAATTGACTTTTATCTGGCAGCTGTCC 203
Dere 151 ----------------------------GATTAT----------------GTAGTGTATGAAATAAA-GGAAAATTGACTTTTATCTGACAGCTGCCC 203
Dana 156 ACACAGCACTCTTGACCTTCGATCACAGTATGTCGCGTATAAATTTGAACGTGAATGGGTGGGTTAAAA-ATAAATATGACTGTTATCTGGCAGCTGTCC 254
Dpse 109 ------------AACATGATTGAAACATTGATTTTATTCCAACCCCAAATCGATGCCAAGACCTTGAAA-ATAAAAATGACTTTTAT-------CTGTCA 188
Dper 109 ------------AACATGATTGAAACATTGATTTTATTCCA-CCCCAAATCGATGCCGAGACCTTGTAAAATAAAAATGACTTTTAT-------CTGTCA 188
Dwil 130 -----------TTGTGTGTTTTAAGATCGATAATTATTGTGGAATTTATGTACTTATAAATACCATGTAGTAAAATAGACTTTTAT-------CTGGCA 210
Dmoj 133 ACCATTTTGCTATCCTAAAAGACGAGTCCCAGCTTATACACAGCGGCCAATTGTTGATTATGAAATAAACGTAAAAGTGACTTTGATACGACAGCTATTT 232
Dgri 129 GTAATACGACATTGAAAGACGAGTACAAATAATAGCCCTATTGCACAAGTTTTGTTAATAAAAGAAAAACCAAAGCTGACTTTAATCTGACAATTGTCA 228
Dvir 163 TTAAACAAAACCCAAAAGACTTGTGCACAAATTTTAAAGCTACAGCTCAAACGTTGATTATTAAATAAAATCACAGATGACTTTGATATGACAGCTGCTC 262

                                                     RCSI
Dmel 207 GCCGTCTATAAGTCTTCTGCG------CCTCTTTTGATGAGCGGCTAATTGGGTTAGC------AAACATCTATAAAGGGCTGGGAATTCGGCCTC---- 290
Dsim 207 GCCGTCTATAAGTCTTCTCCG------CCTCTTCTGATGAGCGGCTAATTGGCTTAGC------AACAATCTATAAAGGGCTGGGAATTCGGCCTC---- 290
Dsec 207 GCCGTCTATAAGACTTCTCCG------CCTCTTTTGATGAGCGGCTAATTGGCTTAGC------AACCATCTATAAAGGGCTGGGAATTCGGCCTC---- 290
Dyak 204 GCCGTCCATTAGCCTTGCCAG-------CCATTTTGATGAACGACTAATTGGCTTAGA------AACCACCTATAAAAGGCCGGAAACGCGGCCTC---- 286
Dere 204 GCCGTCTCCTAGCCTTTCCAG-------CCCTCTTGATGAGCCGCTAATTGGCTTAGG------AACCGCCTATAAAGGGCTGGAAGCTCGGCCTC---- 286
Dana 255 GCTTGTCTCGGTTACGGATTTA------CCAT-------CCCGTCTAATTGGCTTAGA------AAAGGCCTATAAAAGCAGGCCATTCAAGTCGG---G 332
Dpse 189 GTTGGACAGCATTTATTAGCATTAAGACCCCATTTGATAGCTAGCTAATTGGCTTAGA------AACCAGCTATAAAGGTTCCCCGGCCCAGCCG----- 277
Dper 189 GTTGGACAGCATTTATTAGCATTAAGACCCCATTTGATAGCTAGCTAATTGGCTTAGA------AACCAGCTATAAAGGTTCCCCGGCCCAGCCG----- 277
Dwil 211 TCTCGTCAGTGGGCATTAGCATGA--CCCCCCT-----------TAATTGGCTTAGCAGAGTTCAATGGGTATAAAACGGCGGGCAATCGGCGACA--- 293
Dmoj 233 CAAT---------TTATGCATATG----CCA-TCGATCCGCAGCTAATTGGCTCAGA----GCACAGACTATAAAAGCGCTGTCAGTGGCCAGACCG 312
Dgri 229 GATTTGTTCATAGTTTGCATATT----TTA-TTGATCTACGCTTAATTGGCTCAAA---TATTTGCCTATATAAAAGTCACTGTCAACTATTAGC--C 318
Dvir 263 GTTGTACCTGTG-------CCCATT----CCA-TCGATCTGCGCCTAATTGGCTCAAA-----TGCTCGCCTATAAAAGCCACTGTACCATGGCGGCTTC 345

                            Rhodopsin2   exon1                                           Met
Dmel 291 -----------TTGGCACAGATCCACT-TCGATGAGCC-----ATCGTCTAGTCAAAGGGGTTGCGGGTAGCTGAGC-ATG-GAG-CGC-AGT-CAT-TT 367
Dsim 291 -----------TTGGCTCAGATCCACT-TCGATGAGCC-----ATCGTCTAGTCAAAGGGGTTGCGGGTAGCTGAGC-ATG-GAG-CGC-AGT-CAT-TT 367
Dsec 291 -----------TTGGCTCAGATCCACT-TCGATGAGCC-----ATCGTCTAGTCAAAGGGGTTGCGGGTAGCTGAGC-ATG-GAG-CGC-AGT-CAT-TT 367
Dyak 287 -----------TTGGCTCAGATCTAGT-TCGATGAGCC-----ATCGTTTAGTCGAAGGGGTTGCGGGTAGCTGAGC-ATG-GAG-CGC-AGT-CAT-TT 363
Dere 287 -----------TTGGCTCAGATCTAGT-TCGATGAGCC-----ATCGCTTGGTCAAAGGGGCTGCGGGTAGCTGAGC-ATG-GAG-CGC-ACT-CAC-TT 363
Dana 333 CCCTCG-------AGTTCCCACACAGT-TCGATGAGCATTT--GAGACGCTGCACGGCGAGTTGCGGGTAGCTGAGC-ATG-GAG-AGC-ACT-CAT-TT 416
Dpse 278 ------------TGGCTCAGCCACAGT-TCGATGCGC------ATCG-------------GAATCGGGTGACGGAGC-ATG-GAG-CGC-AGT-CTG-TT 339
Dper 278 ------------TGGCTCAGCTACAGT-TCGATGCGC------ATCG-------------GAATCGGGTGACGGAGC-ATG-GAG-CGC-AGT-CTG-TT 339
Dwil 294 -----------TTAGCACGATTTCCACATCAATGTACTTTCAAATC------ACAAA-------CGGATAGCTGAGC-ATG-GAG---------------- 352
Dmoj 313 CACTGC-----TAGTCCGGACTACCTT---GATGTGT---GCTGACCCCCTCGCTAATCGCTGAGCAGCTGCTGAGC-ATG-GAA-CGC-AGC-TTG-TT 395
Dgri 319 AGCTCTA----TTGGCCCGGGTCTCTT---AATGTGTGCCTCCAAATCG----CAAA-----CAGTTAGAACTGAGC-ATG-GTG-CAC-AGT-TTG-AT 396
Dvir 346 AGCTCTGTTAGTTGGCAACCCCTT------GATGTG---CGCCATCGATT---CA-----CGGACGGGTAGCTGAGC-ATG-GAG-----AAG-ACG-TT 419

Dmel 368 G-CCG-GAG-ACG-CCA-TTC-GAC-CTG-GCC-CAT-TCT-----------------GGG-CCC-AGG-TTT-CAG-GCC-CAG-TCG-AGT-GGC-AAT 431
Dsim 368 G-CCG-GTG-ACG-CCA-TTC-GAC-CTG-GCC-CAT-TCT-----------------GGG-CCC-AGG-TTT-CAG-GCC-CAG-TCG-AGC-GGC-AAT 431
Dsec 368 G-CCG-GAG-ACG-CCA-TTC-GAC-CTG-GCC-CAT-TCT-----------------GGG-CCC-AGG-TTT-CAG-GCC-CAG-TCG-AGC-GGC-AAT 431
Dyak 364 G-CCA-GAG-ACG-CCA-TTC-GAC-CTG-GCC-CTT-TCT-----------------GGG-CCC-AGG-TTC-CAG-GCC-CAG-TCG-AGC-GGC-AAT 427
Dere 364 G-CCG-GAG-ACG-CCA-TTG-GAC-TTG-GCC-CTT-TCT-----------------GGG-CCC-AGG-TTT-CAG-GCC-CAG-TCG-AGC-GGC-AAT 427
Dana 417 T-CCG-GAG-ACG-CCA-TTG-ACT-TTG-GCC-CTC-ACA-----------------GGG-CCC-AGG-TTC-CAG-GCC-CAG-TCG-GGC-GGC-AAT 480
Dpse 340 G-CCA-GAG-CCG-CCG-TTG-GCC-ATG-GCA-CTC-CTG-----------------GGG-CCG-AGA-TTC-GAG-GCC-CAA-ACA-GGA-GGC-AAT 403
Dper 340 G-CCA-GAG-CCG-CCG-TTG-GCC-ACG-GCA-CTC-TTG-----------------GGG-CCC-AGG-TTC-GAG-GCC-CAA-ACA-GGA-GGC-AAT 403
Dwil 353 ----------ACG-CCA-TTG-ACC-ATG-GCC-ATG-TCT-----------------GGG-CCC-AGA-TTT-GAG-GCA-CAG-TCG-GGT-GCC-AAT 409
Dmoj 396 G-CCG-CAG-ACA-TTG-GAC-GCT-GCC-TGG-CCT-GTC-AGT-GGA-CTT-GGG-CCC-AGA-TTC-GAT-GCC-CAG-GCG-AGT-GGC-AAT 468
Dgri 397 G-CCA-GAG-ACA-ATC-GCG-GTT-GCC-ATG-TTT-GGC-ACT-CAA-CTT-GGG-CCG-CGA-TTC-GAG-GCC-CAG-GCG-GGC-GGT-AAT 469
Dvir 420 C-GCC-GTT-GCC-ATA-TCG-GGC-TCT-----------------GGA-CTT-GGA-CCA-AGA-TTT-GAG-GCC-CAG-GCG-GGC-CAA-AAT 480
```

**Figure S6**

| Bloomingtons stock number | Gene | Gene symbol | Interactions |
|---|---|---|---|
| 51167 | Abdominal B | Abd-B | Physical |
| 57718 | | CG31612 | Physical |
| 40824 | | CG5446 | Physical |
| 50792 | Deformed | Dfd | Physical |
| 26751 | Deformed | Dfd | Physical |
| 65129 | Enhancer of split mδ, helix-loop-helix | E(spl)mδ-HLH | Physical |
| 41682 | Ets at 65A | Ets65A | Physical |
| 33652 | Inverted repeat binding protein 18 kDa | Irbp18 | Physical |
| 54852 | Poly-glutamine tract binding protein 1 | PQBP1 | Physical |
| 50662 | Sex combs reduced | Scr | Physical, genetic |
| 53991 | Sox21a | Sox21a | Physical |
| 31902 | Sox21a | Sox21a | Physical |
| 34993 | Ultrabithorax | Ubx | Physical |
| 35644 | Abdominal A | Abd-A | Physical, genetic |
| 62873 | chronologically inappropriate morphogenesis | chinmo | Physical |
| 33638 | chronologically inappropriate morphogenesis | chinmo | Physical |
| 35645 | doublesex | dsx | Physical |
| 55646 | doublesex | dsx | Physical |
| 32486 | eyeless | ey | Physical, genetic |
| 65132 | labial | lab | Physical |
| 35413 | ovo | ovo | Physical |
| 42627 | ovo | ovo | Physical |
| 35812 | tiptop | tio | Physical |
| 34067 | yorkie | yki | Physical |
| 37519 | B52 | B52 | Physical |
| 64926 | Antennapedia | Antp | Genetic |
| 32863 | cap-n-collar | cnc | Genetic |
| 36092 | Cyclin E | CycE | Genetic |
| 38902 | Cyclin E | CycE | Genetic |
| 29337 | Distal-less | Dll | Genetic |
| 26752 | engrailed | en | Genetic |
| 33715 | engrailed | en | Genetic |
| 33761 | fushi tarazu | ftz | Genetic |
| 41699 | rhomboid | Rho | Genetic |
| 38199 | scribble | scrib | Genetic |
| 40849 | Calmodulin-binding transcription activator | camta | Mishra et al., 2016 |
| 26225 | defective proventriculus | dve | Mishra et al., 2016 |
| 36721 | longitudenals lacking | lola | Mishra et al., 2016 |
